# UNC-16/JIP3 and UNC-76/FEZ1 limit the density of mitochondria in *C. elegans* neurons by maintaining the balance of anterograde and retrograde mitochondrial transport

**DOI:** 10.1101/279372

**Authors:** Guruprasada Reddy Sure, Anusheela Chatterjee, Nikhil Mishra, Vidur Sabharwal, Swathi Devireddy, Anjali Awasthi, Swetha Mohan, Sandhya P. Koushika

## Abstract

We investigate the role of axonal transport in regulating neuronal mitochondrial density. We show that the density of mitochondria in the touch receptor neuron (TRN) of adult *Caenorhabditis elegans* is constant. Mitochondrial density and transport are controlled both by the Kinesin heavy chain and the Dynein-Dynactin complex. However, unlike in other models, the presence of mitochondria in *C. elegans* TRNs depends on Kinesin light chain as well. Mutants in the three *C. elegans miro* genes do not alter mitochondrial density in the TRNs. Mutants in the Kinesin-1 associated proteins, UNC-16/JIP3 and UNC-76/FEZ1, show increased mitochondrial density and also have elevated levels of both the Kinesin Heavy and Light Chains in neurons. Genetic analyses suggest that, the increased mitochondrial density at the distal end of the neuronal process in *unc-16* and *unc-76* depends partly on Dynein. We observe a net anterograde bias in the ratio of anterograde to retrograde mitochondrial flux in the neuronal processes of *unc-16* and *unc-76*, likely due to both increased Kinesin-1 and decreased Dynein in the neuronal processes. Our study shows that UNC-16 and UNC-76 indirectly limit mitochondrial density in the neuronal process maintaining a balance in anterograde and retrograde mitochondrial axonal transport.

## Introduction

Mitochondria are present along the entire length of the neuron and at greater density in regions with high energy needs such as at the Nodes of Ranvier and at synapses ^1,2^. Additionally, neurons have defined mitochondrial density along neuronal processes that is altered in models of neurodegenerative disease ^3,4^. Majority of the mitochondrial biogenesis in neurons is believed to occur in the cell body ^5,6^. Therefore, transport is likely an important means to control both the presence and density of mitochondria along the neuronal processes. Kinesin-1 and Dynein are the major anterograde and retrograde motors for mitochondrial transport ^7-10^. Kinesin heavy chain in both *Drosophila* and vertebrates is recruited onto the mitochondrial surface via the adaptor Milton/TRAK, in addition, Miro, a Rho GTPase, is essential for mitochondrial transport ^11-14^. Kinesin heavy chain (KHC), independent of Kinesin light chain (KLC), binds to Milton/TRAK ^11,14^. Milton/TRAK in turn binds to Miro present on the mitochondrial surface, this binding facilitates mitochondrial transport ^11,14^. Additionally, RIC-7, a *C. elegans* specific mitochondrial protein, has been shown to control the presence of mitochondria in the motor neuron process ^10^, although its interaction with Kinesin-1 is not established.

Kinesin-1 has multiple interacting proteins, some of which are used to bind cargo ^1,9,11,12,15-20^. UNC-16/JIP3/dSYD scaffolds the JNK pathway kinases and is a major Kinesin-1 adaptor ^15,16,21^. JIP3/UNC-16 facilitates Kinesin-1 mediated transport of activated JNK Kinases ^22^ and Dynein Light Intermediate Chain ^23^. *C. elegans* JIP3/UNC-16 mutants also show an increase in levels of mitochondria, marked by TOMM20, in motor neuron axons ^24^. JIP3/UNC-16 binds to both Kinesin heavy and light chains ^25,26^ and increases motor velocity and run length ^25^. UNC-76/FEZ1 (fasciculation and elongation protein zeta 1) binds to the Kinesin heavy chain subunit and activates the motor ^27-29^.

In this study, we use *C. elegans* touch receptor neuron (TRN) as a model to study mitochondrial transport *in vivo*. The TRN allows us to track mitochondria in a single isolated neuron where its distribution and transport can be easily observed. In wild type animals, mitochondrial density is constant in the TRN. This density is regulated by the Kinesin heavy chain, Kinesin light chains and the Dynein-Dynactin complex. Our data show that the adaptors of Kinesin-1 limit the density of mitochondria in the TRNs by maintaining the balance of anterograde and retrograde mitochondrial transport.

## Results

### Kinesin heavy chain and light chains and the Dynein-Dynactin complex maintain the density of mitochondria along the neuronal process

We visualized mitochondria in the anterior and posterior TRN processes using *jsIs609* ^30^, a transgenic animal expressing matrix targeted GFP in the touch receptor neurons of *C. elegans.* The mitochondrial density in the anterior TRN (ALM anterior lateral mechanosensory) processes in 1-day adult animals is approximately 5 mitochondria/100µm (Figure 1A), similar to those reported earlier ^31^.

**Figure 1:**
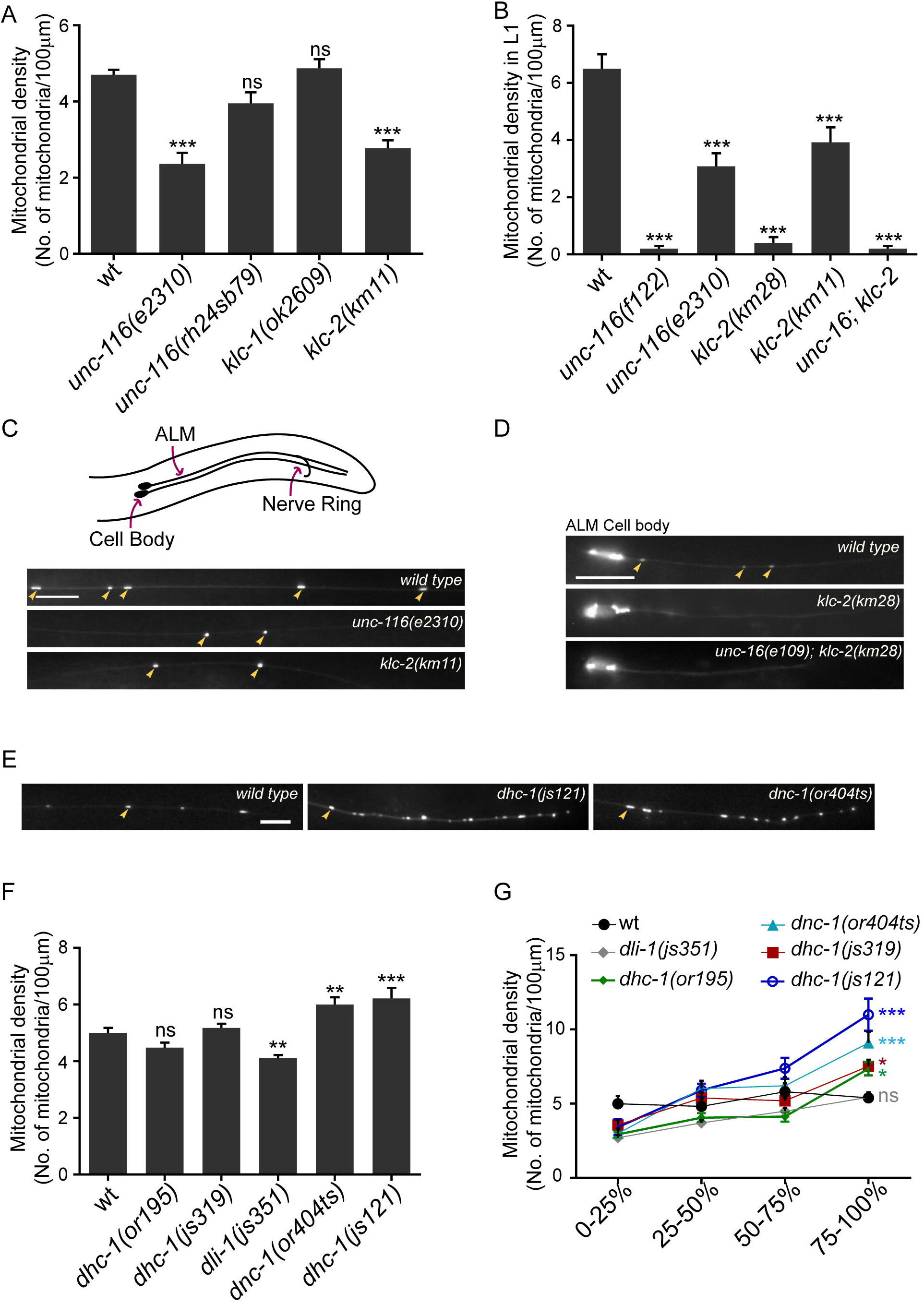
Kinesin-1 and Dynein regulate mitochondrial density in the anterior touch receptor neuronal process. **(A)** Mitochondrial densities in one day adult ALM (anterior lateral mechanosensory) TRNs (touch receptor neurons) of wild type (*jsIs609*) and Kinesin-1 motor subunit hypomorphic mutants. n=16-20 worms. All comparisons made to wild type. **(B)** Average mitochondrial densities in ALM TRN processes of just hatched L1 animals of the following genotypes: Kinesin-1 subunit null alleles (*f122, km28)*, Kinesin-1 subunit hypomorphic alleles (*e2310, km11*) and a double mutant *unc-16(e109); klc-2(km28)*. n=16-20 worms. All comparisons made to wild type. **(C)** Representative images of mitochondria in the ALM TRN processes of wild type (*jsIs609*), *unc-116(e2310)* and *klc-2(km11)* one day adults. Orientation of the anterior TRNs (here only ALM) and schematic representation of the worm is shown above fluorescent images. Scale bar=10μm. **(D)** Representative images of the ALM TRN processes of wild type (*jsIs609*), *klc-2(km28)* and *unc-16(e109); klc-2(km28)* L1 animals. Scale bar=10μm. **(E)** Representative images of the distal ALM TRN processes in wild type (*jsIs609*), *dhc-1(js121)* and *dnc-1(or404ts)* (grown at 22°C). Scale bar=10μm **(F)** Average mitochondrial densities in the ALM TRN processes of Dynein-Dynactin complex mutants. n=10-20 worms. All comparisons made to wild type **(G)** Mitochondrial density across different segments along the ALM TRN processes in wild type, Dynein mutants and Dynactin mutants. n=10-16 worms. All comparisons made to wild type. Colours correspond to respective genotypes. Some mitochondria marked with yellow arrows in C, D and E. Data represented as Mean ± SEM. Statistical tests used for A, B and F: One-way ANOVA, for G: Two-way ANOVA. All tests use Bonferroni multiple comparisons correction, ns: not significant, p value *<0.05, **<0.01, ***<0.001.

To investigate the role of motors, we examined mitochondrial density in the loss of function mutants of the UNC-116/Kinesin Heavy Chain (KHC) and the two Kinesin Light Chain genes, k*lc-1* and *klc-2*. Partial loss of function alleles *unc-116*(*e2310)*^32^ and *klc-2(km11*)^21^ show significant reductions in their mitochondrial densities in the anterior TRN processes in both adult and L1 animals (Figure 1A, B, C). *unc-116(rh24sb79)*^33^ has been shown to reduce mitochondrial density in motor neurons^10^ although we do not see a significant decrease in the anterior TRN (Figure 1A). The null alleles *unc-116(f122)*^34^ and *klc-2(km28)*^21^ show a complete absence of mitochondria along the neuronal process in just hatched L1 animals compared to similarly staged wild type L1 (Figure 1B, D). We could not examine the adult null mutants of the Kinesin-1 complex since they do not survive beyond the early L2 stage. The mitochondrial density in *klc-1*(*ok2609*)^35^ with a 1383 bp deletion in the gene was similar to that observed in wild type (Figure 1A). Thus, both Kinesin-1 motor subunits UNC-116/KHC and KLC-2, but not KLC-1 play an essential role in maintaining mitochondrial density in the TRN process.

Dynein is known to transport mitochondria in the retrograde direction ^7^. Thus, we assessed mitochondrial density in the mutants of the Dynein heavy chain (*dhc-1*), Dynein light intermediate chain (*dli-1*) and the p150 Glued Dynactin subunit (*dnc-1*). We observed that a strong loss-of-function allele, *dhc-1(js121)*^36^, and *dnc-1(or404ts)*^36^ grown at the restrictive temperature, show a significant increase in overall mitochondrial density (Figure 1F). All other *dhc-1* mutant alleles examined show average overall densities of mitochondria, similar to wild type (Figure 1F). However, all Dynein-Dynactin complex mutants, except *dli-1(js351)*^36^, show a significant increase in mitochondrial density at the distal quarter of the neuronal process (Figure 1E, G). This phenotype is similar to that reported in Dynein complex mutants, where Dynein-dependent cargo accumulate at the end of the anterior TRN process ^36^. Additionally, *dhc-1(js121), dhc-1(or195)*^37^ and *dnc-1(or404ts)* all show a significant increase compared to wild type in the ratio of density of mitochondria in the last quarter of the anterior TRN to the first quarter (Figure S1A). Despite a reduction in average overall mitochondrial density (Figure 1F), *dli-1(js351)*, shows nearly a doubling of but not a statistically significant increase compared to wild type in the ratio of mitochondrial density in the distal TRN process to the proximal TRN process (Figure S1A). The *dli-1(js351)* fold change in mitochondrial density between the last quarter and first quarter of the neuronal process is similar to that observed in the hypomorphic *dhc-1(js319)* allele (Figure S1A). *dhc-1(js319)* that shows a significant accumulation of mitochondria in the distal anterior TRN (Figure 1G). *dli-1(js351)* animals are sterile and are the progeny of *dli-1/+* hermaphrodites ^36^. The weaker phenotype observed in *dli-1(js351)* animals thus may arise from the known maternal perdurance of wild type DLI-1 ^38^.

Mitochondrial transport in both mammalian cells and *Drosophila* neurons has been shown to occur independent of the Kinesin Light Chain ^14^. The dependence of mitochondrial density on KLC-2 suggests that *C. elegans* may have alternate and/or additional means to regulate mitochondrial transport in neurons, for instance, through RIC-7 ^10^. The complete absence of mitochondria in the null mutants of *klc-2* suggests that KLC-2 in conjunction with UNC-116 is an essential complex that mediates mitochondria entry into axons of *C. elegans.* The observed distal accumulation of mitochondria in Dynein and Dynactin mutants suggests a role for the Dynein-Dynactin complex in facilitating retrograde mitochondrial transport in *C. elegans* TRNs.

### *miro* mutants do not significantly alter TRN mitochondrial density

The Miro-Milton complex is the major mitochondrial adaptor for Kinesin-1 ^13,14^. No Milton homologues have been found in the *C. elegans* genome ^39^, but three Miro orthologues have been reported to be present ^40^. Therefore, we investigated the role of all reported *miro* genes, whose mutants lack the mitochondria associated transmembrane domain, some of their EF hands and GTPase domains (Figure S2). *C. elegans* MIRO-1 shows ∼60% protein sequence similarity with *D. melanogaster* Miro while *C. elegans* MIRO-2 and MIRO-3 show ∼30% protein sequence similarities ^41^. *miro-1(tm1966), miro-2(tm2933)* and *miro-3(tm3150)* single mutants do not alter mitochondrial density in the anterior TRN process (Figure 2A). Double mutants [*miro-1; miro-2*], [*miro-1; miro-3*] and [*miro-3; miro-2*] do not show a significant change in mitochondrial density from wild type. Only *miro-3* mutants show significantly lower density compared to all three double mutants [*miro-1; miro-2*], [*miro-1; miro-3*] and [*miro-3; miro-2*] in the neuronal process (Figure 2A).

**Figure 2:**
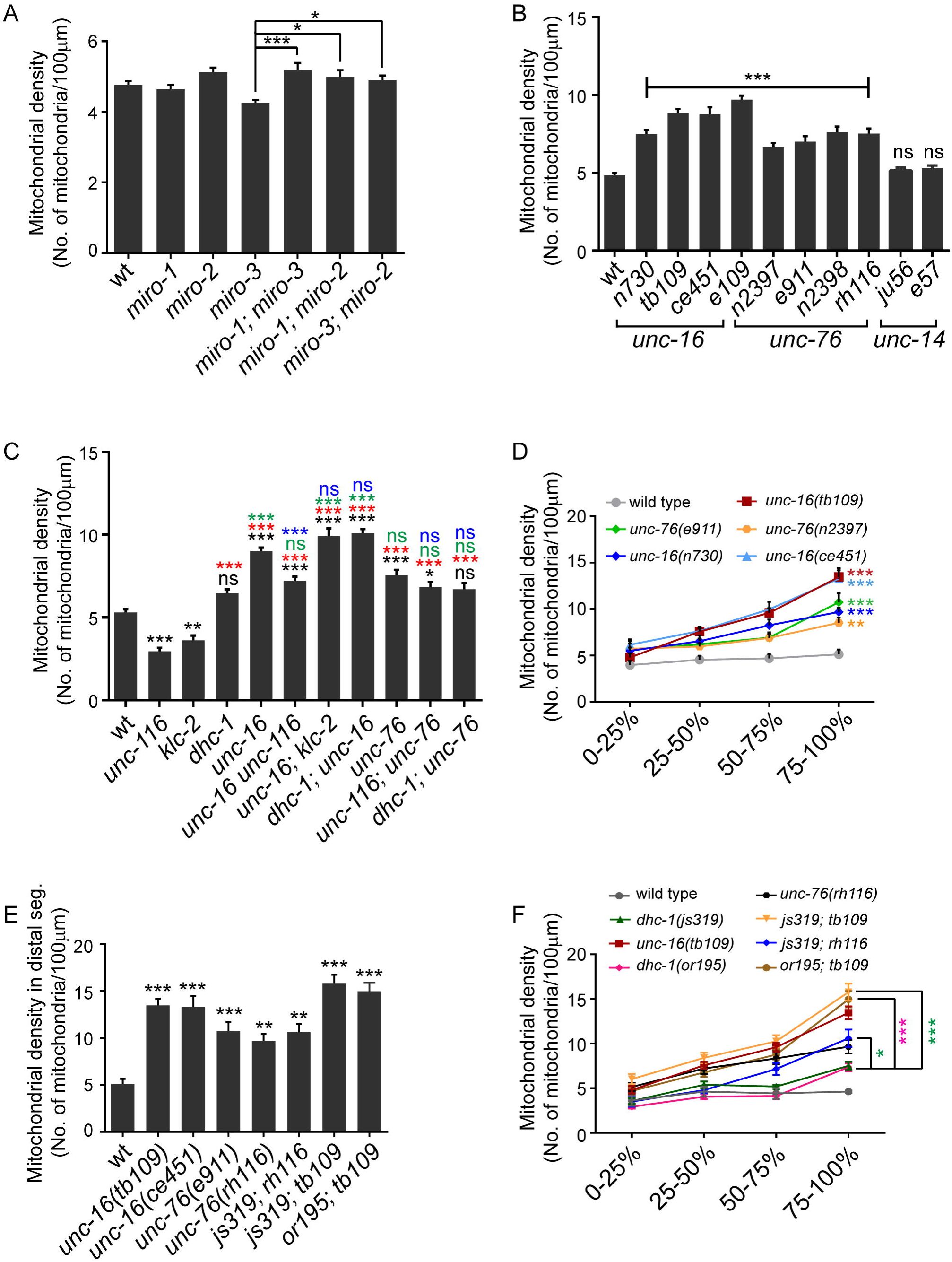
Kinesin-1 associated proteins- UNC-16/JIP3 and UNC-76/FEZ1- regulate mitochondrial density in the anterior touch neuronal process. **(A)** Mitochondrial density in the ALM (anterior lateral mechanosensory) touch neuronal processes of one day adult wild type (*jsIs609*), *miro-1(tm1966), miro-2(tm2933), miro-3(tm3150)* and double mutants [*miro-1(tm1966); miro-3(tm3150)*], [*miro-1(tm1966); miro-2(tm2933)*] and [*miro-3(tm3150); miro-2(tm2933)*]. n≥22 worms. Only significant comparisons have been marked. (**B)** Mitochondrial density in one day adult ALM neuronal processes of mutants in other Kinesin-1 associated proteins-*unc-16(n730), unc-16(tb109), unc-16(ce451), unc-16(e109), unc-76(n2397), unc-76(e911), unc-76(n2398), unc-76(rh116), unc-14(ju56)* and *unc-14(e57).* n≥10 worms. All comparisons made to wild type (*jsIs609*). **(C)** Mitochondrial densities in one day adult wild type (*jsIs609*), *unc-116(e2310), klc-2(km11), dhc-1(js319), unc-16(tb109), unc-76(rh116)* and double mutants between Kinesin-1 adaptors and motor hypomorphs [*(tb109 e2281), (tb109; km11), (js319; tb109), (e2310; rh116)* and *(js319; rh116)*]. *e2310* and *e2281* are the same allele, see Table S1. n≥18 worms. Black * or ns (not significant) are comparisons to wild type. Green * or ns are comparisons of adaptor single mutants and double mutants with adaptors to *dhc-1* single mutant. Red * or ns are comparisons of *dhc-1*, adaptor single mutants and double mutants with adaptors to both *unc-116* and *klc-2*. Blue * or ns are comparisons of motor subunit; adaptor double mutants to respective adaptor single mutants. **(D)** Mitochondrial density across different segments along the neuronal process in one day adult wild type (*jsIs609*) and mutants in Kinesin-1 associated proteins. n≥18 worms. All comparisons made to wild type. Colours correspond to respective genotypes. **(E)** Mitochondrial density in the distal segment (last quarter) in wild type (*jsIs609*), *unc-16(tb109), unc-16(ce451), unc-76(e911), unc-76(rh116)*, [*dhc-1(js319); unc-76(rh116)*], [*dhc-1(js319); unc-16(tb109)*] and [*dhc-1(or195); unc-16(tb109)*]. n≥10 worms. All comparisons made to wild type. **(F)** Mitochondrial densities in one day adults of wild type (*jsIs609*), Dynein mutants, *unc-16, unc-76* and double mutants between Kinesin-1 binding proteins and Dynein mutants across different segments along the neuronal process. n≥ 10 worms. All comparisons made between double mutants and the corresponding dynein complex single mutant. Data represented as Mean ± SEM. Statistical tests one-way ANOVA (A, B, C, E) and two-way ANOVA (D, F) with Bonferroni multiple comparisons correction, ns: not significant, p value *<0.05, **<0.01, ***<0.001.

*miro-1* and *miro-2* RNAi has been shown to reduce mitochondrial density and over-expression of MIRO-1 increases mitochondrial density in *C. elegans* GABA neurons ^42^. Additionally, Miro-1 mutants show a reduction in the amount of mitochondria in the hypodermis ^43^. Our results do not rule out a role for Miro in mitochondrial movement along the neuronal process but suggest that the Kinesin Heavy Chain-Miro complex may not be critical for mitochondrial exit from the cell body of touch neurons.

### Kinesin-1 associated proteins UNC-16/JIP3 and UNC-76/FEZ1 regulate mitochondrial density

Since the *miro* genes did not have a significant effect on mitochondrial density, we investigated mutants in other Kinesin-1 associated proteins on mitochondrial density. Three proteins, UNC-16/JIP3, UNC-76/FEZ1 and UNC-14 are known to associate with the Kinesin-1 complex and play multiple roles in transport of cargo ^15,16,21,25,28,29^. Multiple alleles of *unc-16* (*e109*^44^, *tb109*^45^, *n730*^44^ *and ce451*^24^) show a significant increase in mitochondrial density along the anterior TRN process compared to wild type (Figure 2B). *unc-76* alleles (*n2397*^27^, *n2398*^27^, *e911*^27,46^ and *rh116*^27^) show a significant but less pronounced increase in mitochondrial density than *unc-16* when compared to wild type (Figure 2B). However, *unc-14 (e57*^47^ and *ju56*^21^) alleles do not show a significant change in mitochondrial density compared to wild type in the TRN processes (Figure 2B). A significant increase in mitochondrial density is observed in anterior TRNs of L1 larval stage *unc-16(tb109)* and *unc-76(n2397)* animals (Figure S1B).

Since UNC-16 and UNC-76 affect anterograde axonal transport through their interaction with Kinesin-1 ^21,28^, we examined whether the increased mitochondrial density regulation in *unc-16* and *unc-76* depends on the presence of Kinesin-1. Double mutants between *unc-16(e109)* and the null allele *klc-2(km28)* result in an absence of mitochondria from the neuronal process identical to *klc-2(km28)* alone (Figure 1B, D). We were unable to build recombinants between three different *unc-16* alleles and the *unc-116(f122)* null allele perhaps due to a combination of the lethality of *f122* allele and that the two genes are about 0.2 map units apart. Using double mutants with hypomorphic alleles in the Kinesin-1 subunits, we observe that [*unc-16 unc-116*], [*unc-16; klc-2*] and [*unc-116*; *unc-76*] show significantly increased mitochondrial density compared to both wild type (Figure 2C, black*) and the respective Kinesin-1 subunit single mutant alleles (Figure 2C, red*). *unc-16 unc-116* animals show a significant reduction in mitochondrial density compared to *unc-16* alone (Figure 2C, blue*). *unc-16; klc-2* and *unc-116; unc-76* animals do not show a significant reduction in mitochondrial density compared respectively to *unc-16* and *unc-76* alone (Figure 2C, blue ns). In conjunction with *unc-16; klc-2(km28)* (Figure 1B), we conclude that, Kinesin-1 appears necessary for increased mitochondrial density at least in *unc-16*.

Both hypomorphic alleles *unc-116(e2310)* and *klc-2(km11)* produce lower levels of full length UNC-116 and KLC-2 respectively ^48^. One possible explanation to account for the elevated mitochondrial density in double mutants between the adaptors and hypomorphic alleles of Kinesin-1 is that, presence of even low levels of the Kinesin-1 motor maybe sufficient to cause increased mitochondrial density in *unc-16* and *unc-76* mutants.

### *unc-16* and *unc-76* animals have elevated levels of Kinesin-1 in the neuronal process

Hypomorphic mutations in either Kinesin heavy chain or Kinesin light chain genes do not substantially reduce the increased mitochondrial density observed in *unc-16* and *unc-76* mutant animals. Therefore, we sought to determine if Kinesin-1 levels were altered in *unc-16* and *unc-76* mutants. Using transgenic UNC-116::GFP and KLC-2::GFP lines, driven by their endogenous promoters, that rescue their respective null alleles ^48^, we observe an increase in levels of UNC-116::GFP and KLC-2::GFP in neurons of both *unc-16* and *unc-76* animals (Figure 3A, C, E, F). This increase is seen early in development in L1 animals as well (Figure S3A, B). This observation is consistent with a significant overall increase in endogenous protein levels of ∼30% in UNC-116 and ∼300% in KLC-2 in *unc-16(tb109)* mutants (Figures 3H, S3C, S3D). We do not see an increase in Kinesin-1::GFP levels in non-neuronal cells like the seam cells of *unc-16* animals (Figure 3B and 3D) suggesting that the overall increase may arise largely from an increase of motor levels in neurons. KLC-2::GFP but not UNC-116::GFP levels increase in the seam cells and perhaps other non-neuronal cells of *unc-76* animals (Figure 3B, D, S3A, B). Levels of *unc-116* and *klc-2* mRNA in two alleles each of *unc-16* and *unc-76* do not show any change suggesting that the increase in Kinesin-1 motor levels in these mutants may occur through a post-translational step (Figure S3E). Additionally, in *unc-16 unc-116*, there is a 40% increase in overall UNC-116 protein levels (Figure 3H and S3D, p=0.06) compared to *unc-116.* This small increase may partially contribute to the elevated mitochondria density seen in *unc-16 unc-116* animals (Figure 2C).

**Figure 3:**
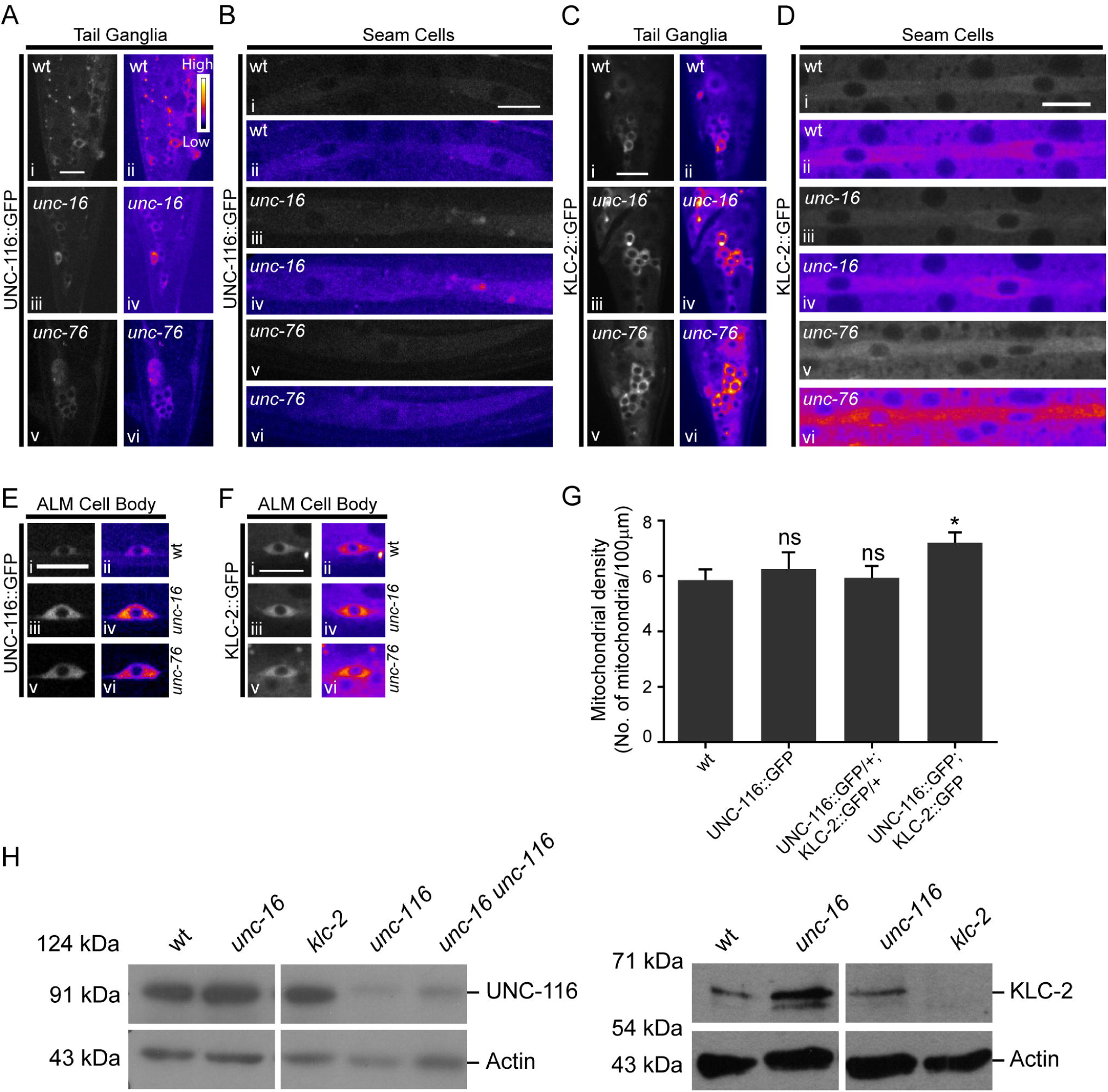
*unc-16* and *unc-76* mutants have elevated levels of Kinesin-1 subunits in neurons. **(A)** Representative images and corresponding heat maps of UNC-116::GFP in wild type, *unc-16(e109)* and *unc-76(n2398)* tail ganglia of L4 animals. **(B)** Representative images and corresponding heat maps of UNC-116::GFP in wild type, *unc-16(e109)* and *unc-76(n2398)* in seam cells of L4 animals. **(C)** Representative images and corresponding heat maps of KLC-2::GFP in wild type, *unc-16(e109)* and *unc-76(rh116)* tail ganglia of L4 animals. **(D)** Representative images and corresponding heat maps of KLC-2::GFP in L4 wild type, *unc-16(e109)* and *unc-76(rh116)* seam cells. **(E, F)** Representative images and corresponding heat maps of UNC-116::GFP (E) and KLC-2::GFP (F) in ALM TRN cell bodies of wild type, *unc-16(e109)* and *unc-76(n2398)* L4 animals. **(G)** Average mitochondrial density in the PLM (posterior lateral mechanosensory) touch neuronal process of L1 when Kinesin-1 subunits are over-expressed. Data represented as Mean ± SEM, n≥8 worms. Mann Whitney test, p value *<0.05. **(H)** Western Blots showing levels of Kinesin-1 protein subunits in whole worm lysates prepared from wild type, *unc-16(tb109)* and *unc-16(tb109) unc-116(e2281)* animals. Lysates prepared from *unc-116(e2310)* and *klc-2(km11)* were used as controls respectively for the UNC-116 and KLC-2 antibodies. *e2310* and *e2281* are the same allele, see Table S1. The experiment was performed at least three independent times in duplicates, quantitation shown in supplementary figure 3D. Scale bar:10μm.

The increase in Kinesin-1 motor levels in *unc-16* and *unc-76* mutant animals may at least partially account for the increase in mitochondrial density in these animals. To test whether an elevation in Kinesin-1 levels alone is sufficient to result in an increase in mitochondrial density in the touch receptor neurons, we measured the density of mitochondria after over-expression of UNC-116::GFP and KLC-2::GFP in wild type animals using *jsIs1073*, with tagRFP targeted to the mitochondrial matrix^49^. Animals over-expressing Kinesin heavy chain alone or wild type animals heterozygous for both Kinesin heavy and light chain-2 expressing transgenes do not show an increase in mitochondrial density (Figure 3G). Homozygous animals over-expressing both UNC-116::GFP and KLC-2::GFP encoding transgenes, show a significant increase in mitochondrial density in the TRN process (Figure 3G). This suggests that elevation of just the Kinesin heavy chain or lower levels of increase in Kinesin-1 may be insufficient to elevate mitochondrial density. The smaller increased mitochondrial density in animals homozygous for the Kinesin heavy and light chain transgenes compared to *unc-16* and *unc-76* may arise from the relatively modest increase in levels of Kinesin-1 in a wild type background compared to the increase in motor levels seen in *unc-16* and *unc-76* animals (Figure 3E, F). Additionally, the elevation of just the anterograde motor levels in *unc-16* and *unc-76* may be insufficient to account for the observed large increase in density of mitochondria in the TRNs of these mutants.

### UNC-16 regulates axonal mitochondrial density in part through Dynein

UNC-16 has a role in retrograde transport as an adaptor of Kinesin-1, by facilitating binding of KLC-2 to Dynein ^23^. Additionally, UNC-76 associates with the Kinesin heavy chain and activates Kinesin-1, thus facilitating the transport of Kinesin-1 cargo, one of which is Dynein ^23,29,50,51^. Thus both mutants are likely to reduce Dynein mediated transport through different mechanisms.

We first examined whether *unc-16* and *unc-76* mutants shared mitochondrial distribution phenotypes with mutants in the Dynein complex. *unc-16* but not *unc-76* has an overall mitochondrial density significantly above that of *dhc-1* (Figure 2C, green*). However, similar to Dynein-Dynactin mutants, both *unc-16* and *unc-76* animals also show a significant increase in mitochondrial density in the distal quarter of the neuronal process (Figure 1G and Figure 2D). In all alleles of both mutants examined, the density of mitochondria in the last quarter of the TRN process is significantly higher than that observed in wild type (Figure 2D, E). The distal accumulation phenotype is more severe in the strong loss of function *unc-16* alleles^24,45^ (*ce451, tb109)* compared to the strong loss of function *unc-76* alleles^27^ (*rh116, n2397, e911*) (Figure 2D, E). The phenotypes where mitochondria accumulate towards the end of the neuronal process are consistent with the known accumulation of Dynein-dependent cargo at the distal end of the anterior TRN ^36^. The distal mitochondrial accumulation phenotypes in *unc-16* and *unc-76* may be accounted for by mitochondria moving to the distal part of the neuronal process using anterograde motors and accumulating towards the end of the neuronal process due to their inability to return to the cell body due to insufficient functional Dynein activity. We observe that two alleles of both *unc-16* and *unc-76* have *dhc-1* mRNA levels similar to wild type (Figure S3E). We were unable to effectively image DHC-1 in neurons as the *dhc-1* promoter driven DHC-1::GFP line was very faint ^52^.

Since, both UNC-16 and UNC-76 contribute to the anterograde transport of Dynein, we investigated the genetic interactions between mutants in both genes with a Dynein complex mutant. We examined whether a further decrease in Dynein function exacerbated the distal TRN mitochondria accumulation phenotypes of *unc-16* and *unc-76*.

[*dhc-1; unc-16*] and [*dhc-1; unc-76*] do not show any significant increase in overall mitochondrial density compared respectively to *unc-16* and *unc-76* alone (Figure 2C, blue ns). Further the overall mitochondria density in *dhc-1; unc-16* but not in *dhc-1; unc-76* is significantly higher than the *dhc-1* single mutant (Figure 2C, green* & ns). Double mutants [*dhc-1(js319); unc-16(tb109)*], [*dhc-1(or195); unc-16(tb109)*] and [*dhc-1(js319); unc-76(rh116)*] all continue to show greater numbers of distal mitochondria compared to wild type (Figure 2E). The distal mitochondrial density in all three double mutants listed above are significantly higher compared to the respective dynein heavy chain single mutants (Figure 2F). [*dhc-1(js319); unc-16(tb109)*], [*dhc-1(or195); unc-16(tb109)*] and [*dhc-1(js319); unc-76(rh116)*] show a small but a statistically not significant increase in the distal mitochondrial density (17%, 11% and 10% respectively) compared to the distal mitochondria accumulation seen in the corresponding the *unc-16* or *unc-76* single mutant alleles in the same region (Figure 2F). The lack of a bigger increase could arise from the combination of alleles used i.e. weak loss of function mutants in the Dynein complex built with strong loss of function alleles of *unc-16* and *unc-76*.

We think that reduced Dynein transport could contribute to the increased density of mitochondria at the distal end of the neuron in *unc-16* and *unc-76* animals. In *unc-16* animals, due to the absence of the adaptor, anterograde transport of Dynein and retrograde transport of Dynein-dependent vesicles has been shown to be reduced ^23^, this may account for the more prominent mitochondrial distal accumulation in this adaptor mutant. The absence of UNC-76/FEZ1 may lead to weaker activation of the Kinesin-1 motor, also potentially reducing the available Dynein at the end of the neuronal process. However, other molecules such as Kinesin light chain are also known to activate Kinesin-1 ^53,54^. The redundancy in Kinesin-1 activation along with elevated levels of Kinesin-1 seen in *unc-76* may lead to greater levels of Dynein in the end of the neuron in *unc-76* compared to *unc-16*; perhaps accounting for the less severe distal mitochondrial accumulation phenotype in *unc-76* compared to *unc-16.*

### *unc-16* and *unc-76* show increased anterograde flux when Kinesin-1 levels are limiting

Our genetic data suggests that the altered mitochondrial density in these adaptor mutants may arise indirectly due to an altered ratio of anterograde and retrograde motors leading to changes in transport. Thus, we examined mitochondrial transport in *unc-16* and *unc-76* mutants.

Our data show that anterograde mitochondrial movement is dependent on the Kinesin-1 complex since mutants in both *unc-116(e2310)* and *klc-2(km11)* have significantly reduced anterograde flux compared to wild type animals (Figure 4A, black*). Surprisingly, *dhc-1* mutants also show a significant (∼68%) decrease in anterograde transport compared to wild type (Figure 4A, black*). Despite an increase in Kinesin-1 motor levels both alleles of *unc-16* and *unc-76(n2397)* do not show a significant increase in anterograde transport compared to wild type (Figure 4A). Puzzlingly, *unc-76*(*rh116)*, shows a significant reduction in the anterograde flux compared to wild type, contrary to the expectation that increased Kinesin-1 motors should lead to increased anterograde flux (Figure 3D, F, Figure 4A). However, the anterograde mitochondrial flux in *unc-16 unc-116* is significantly increased compared to *unc-116* (red*) perhaps arising from the 40% elevation (p=0.06) in anterograde motor levels seen in the double mutant compared to *unc-116* alone (Figure 3H, S3D). *unc-16 unc-116* has significantly lower anterograde flux compared both to wild type (black*) and the *unc-16* single mutant (blue*) (Figure 4A). This might arise due to the reduction in UNC-116 levels in *unc-16 unc-116* compared to wild type and *unc-16* alone (Figure 3H). Double mutants [*unc-16 unc-116*], [*unc-16; klc-2*] and [*unc-116; unc-76*] show at least a 250% increase in anterograde mitochondrial flux, compared to the corresponding Kinesin-1 single mutants alone (Figure 4A, red*). This suggests a recovery of anterograde mitochondrial flux may occur in part due to an increase in Kinesin-1 levels in these double mutants. These data support the hypothesis that the lack of UNC-16 and UNC-76 can limit anterograde transport of mitochondria in the neuronal processes at least when Kinesin-1 levels are significantly reduced. However a simple explanation that there is an increase in Kinesin-1-dependent anterograde transport probably mediated through the elevated levels of UNC-116 and KLC-2, thereby contributing to increased mitochondrial density in *unc-16* and *unc-76*, may be insufficient.

**Figure 4:**
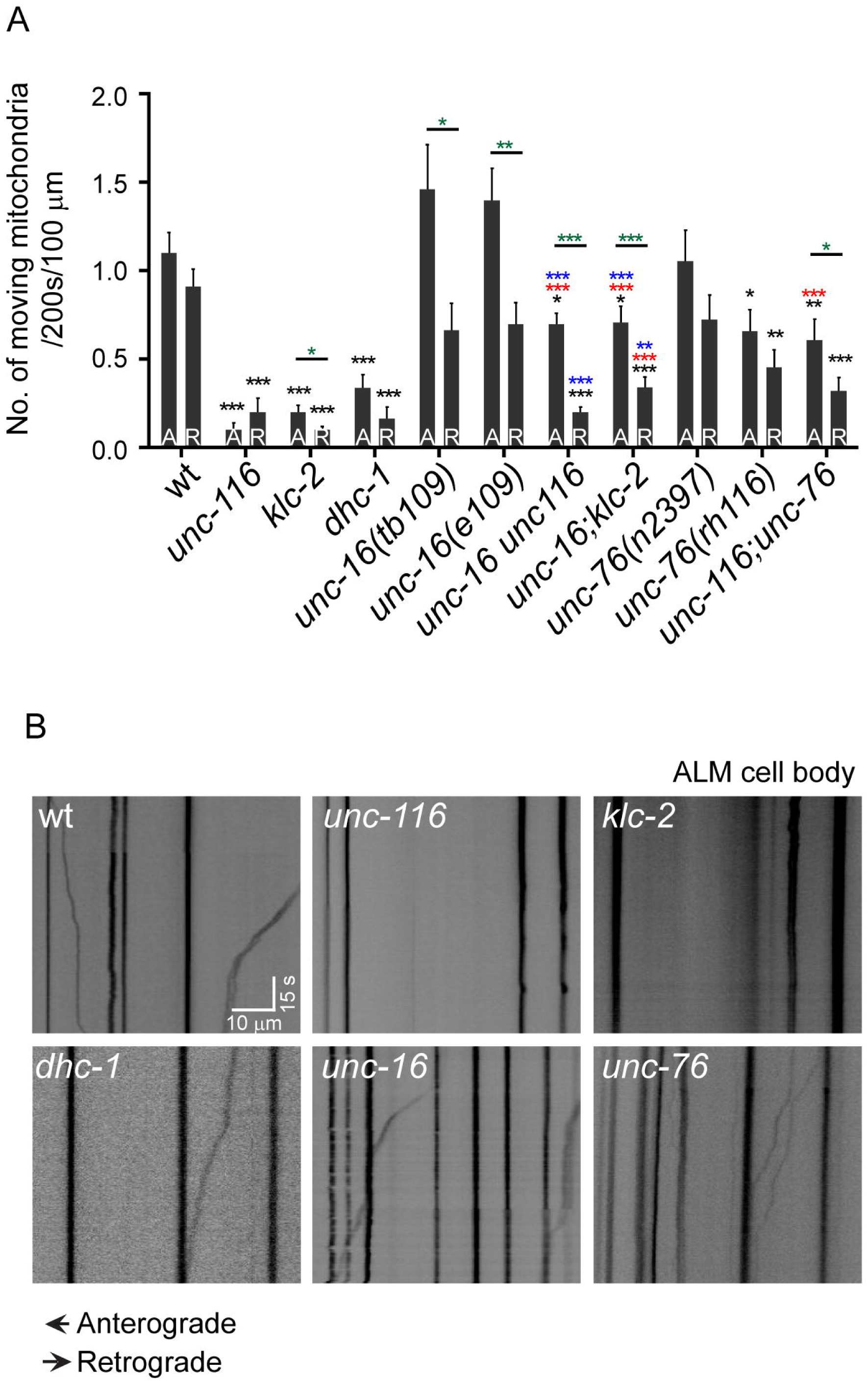
UNC-16 and UNC-76 change mitochondrial flux in anterior TRNs. Mitochondrial anterograde and retrograde flux in the distal ALM processes in L3 animals of **(A)** wild type (*jsIs609*), *unc-116(e2310), klc-2(km11), dhc-1(js319), unc-16(tb109), unc-16(e109), unc-76(n2397), unc-76(rh116)* and double mutants [*unc-16(e109) unc-116(e2310)*], [*unc-16(e109); klc-2(km11)*] and [*unc-116(e2310); unc-76(rh116)*]. Data represented as Mean ± SEM. A: Anterograde; R: Retrograde. n ≥ 34 worms. Statistical tests used: The above multiple comparisons were done using multiple t-test with Sidak-Bonferroni correction. Black * are comparisons to respective wild type direction of movement. Red * are comparisons to respective directions of movement between motor; adaptor double mutants and corresponding motor subunit single mutants. Blue * are comparisons to respective directions of movement between motor; adaptor double mutants and corresponding adaptor single mutants. Green * are comparisons between anterograde and retrograde movements in the same genotype. p value *<0.05, **<0.01, ***<0.001. Comparisons that are not marked eg retrograde transport in *tb109* to retrograde transport in wild type, are not statistically significant. **(B)** Representative kymographs of L3 wild type, *unc-116(e2310), klc-2(km11), dhc-1(js319), unc-16(tb109)* and *unc-76(n2397)* animals. Scale bars: x axis 10 μm, y axis 15 seconds.

### *unc-16* and *unc-76* mutants show an altered ratio of anterograde to retrograde transport

We examined if changes in retrograde transport could account for the elevated mitochondrial density in *unc-16* and *unc-76*. We observed significantly reduced retrograde flux of mitochondria in *unc-76(rh116), dhc-1, klc-2* and *unc-116* animals compared to wild type (Figure 4A black*, B). Retrograde mitochondrial flux is reduced by ∼82% in *dhc-1* animals compared to wild type. The reduced retrograde flux in *unc-116(e2310)* and *klc-2(km11)* likely arises from both the fewer mitochondria in the neuronal process available for retrograde transport and the known role of these motors in transporting Dynein in the axon ^23^. However neither allele of *unc-16* nor *unc-76(n2397)* show significant changes in retrograde mitochondrial flux compared to wild type (Figure 4A), despite a demonstrated reduction in Dynein transport at least in *unc-16* ^23^. The double mutants [*unc-116 unc-16*], [*unc-16; klc-2*] and [*unc-116; unc-76*] show at least a 60% significant reduction in retrograde mitochondrial flux, compared to wild type (Figure 4A, black*). The double mutants [*unc-116 unc-16*], [*unc-16; klc-2*] have significantly lower retrograde transport compared to *unc-16* alone, consistent with a reduced amount of Dynein transport (Figure 4A, blue*). *unc-116; unc-76* does not show a significant reduction in retrograde mitochondrial transport compared to *unc-76* alone, perhaps as there may not be as strong an effect of lack of UNC-76 on Dynein transport. The changes in retrograde transport are complex and not easily accounted for by a simple model where all Dynein is transported with the help of UNC-16 and UNC-76 regulated pathways. Mere changes in retrograde transport cannot readily account for the observed increase in mitochondrial density in *unc-16* and *unc-76*.

The net mitochondrial density probably arises due to a balance in transport of anterograde and retrograde flux, thus we decided to examine the ratios of mitochondrial transport in the different mutants. Although there is little statistical change in the anterograde flux of *unc-16(tb109)* and *unc-16(e109)* animals, there is an increase of >30% compared to wild type (Figure 4A). *unc-76(n2397)* animals show a 10% increase and *unc-76(rh116)* animals show a 38% decrease in anterograde flux respectively compared to wild type. *unc-16(tb109)* and *unc-16(e109)* animals respectively show a 13% and 22% reduction in retrograde flux compared to wild type. *unc-76(n2397)* and *unc-76(rh116)* animals respectively show a 16% and 50% decrease in retrograde mitochondrial flux when compared to wild type. We thus examined whether the difference in anterograde and retrograde transport or its ratio might account for the observed overall mitochondrial density increase observed in adaptor mutants and adaptor; motor subunit double mutants. Anterograde and retrograde flux of mitochondria do not differ from each other in wild type but significantly differ from each other in both *unc-16* mutants and all adaptor; Kinesin-1 subunit double mutant combinations examined (Figure 4A, green*). These genetic backgrounds all significantly elevate mitochondrial density. Notably, while the anterograde and retrograde flux of mitochondria balance each other in wild type animals, mutants in *unc-16(tb109), unc-16(e109), unc-76(n2397), unc-76(rh116)* and *dhc-1(js319)* have higher anterograde/retrograde flux ratios respectively of 1.9, 2, 1.6, 1.5 and 2 compared to the wild type ratio of ∼1. The double mutants [*unc-116 unc-16*], [*unc-16; klc-2*] and [*unc-116; unc-76*] have an anterograde to retrograde flux ratio of 3.5, 2.1 and 1.9 respectively. All genotypes that show an anterograde/retrograde flux ratio much greater than 1 also have significantly increased mitochondrial density in the neuron compared to wild type (flux ratio of ∼1). These data suggest that, UNC-16 and UNC-76 may act to maintain the balance between anterograde and retrograde flux of mitochondria in the neuronal processes. This anterograde bias in the ratio of transport in *unc-16* and *unc-76* may contribute in part to the increased mitochondrial density in the TRN processes of these genotypes.

## Discussion

The results described in this study suggest that, the Kinesin-1 mediated regulation of mitochondrial transport by UNC-16 and UNC-76 could occur at two levels. (i) The increase in Kinesin-1 levels in adaptor mutants. This increase coupled with the inability of the increased Kinesin-1 to recruit UNC-16 and UNC-76 dependent cargo may lead to more Kinesin available for mitochondrial anterograde transport. This is consistent with the striking observation where double mutants between Kinesin-1 and the adaptor mutants show higher levels of anterograde mitochondrial transport compared to motor mutants alone. (ii) In addition to the anterograde transport effects, we also see reduced retrograde transport. This may be due to reduced availability of dynein complex subunits at the distal end of the neuron as has been reported ^23^. Additionally, UNC-16 is also independently known to interact with the retrograde transport complex ^55^. These dual effects may both contribute to the increased density of mitochondria in the neuronal process with the greatest density at the distal end of the process.

### Mitochondrial density regulation in neurons is a complex process

Cultured neurons from rats and zebrafish sensory neurons *in vivo* show a defined density of mitochondria ^56,57^. This mitochondrial density changes, both, during development ^57^ and when neurons express human disease causing variants of TDP-43 and SOD1 ^56^. We show that wild type *C. elegans* TRNs also have a defined density of mitochondria. Localization of mitochondria in specific regions in the neuronal process and their overall density in the axon are likely dependent on multiple factors. In our study, we find that changes in axonal transport act as one factor in regulating density of mitochondria. However, there are likely to be additional factors. This is illustrated by the vast reduction in anterograde transport in a weaker loss-of-function allele *dhc-1* mutant, perhaps as a compensatory mechanism to maintain overall density of mitochondria in the neuronal process. Thus, we think that the relationship between motor levels, mitochondrial flux and average axonal mitochondrial density is likely to be complex.

## Materials and Methods

### Worm maintenance and imaging

Worms were grown using standard protocols ^58^, mitochondrial density was assessed in worms grown without any contamination using 30 mM sodium azide or 10 mM sodium azide as an anesthetic. Each animal was imaged within 10 minutes of anesthetisation. Overlapping images were used to reconstruct the entire neuronal process. The neuronal length was measured using the background fluoresecence in the TRN and the total number of mitochondria were counted. Mitochondrial density was calculated as number of mitochondria/100 micron of the neuronal process using 20-25 animals. L3 animals were anaesthetized in 3 mM Levamisole solution and time lapse images were acquired at 2fps for 200s in the distal regions of the ALM (Anterior Lateral Mechanosensory neuron listed as anterior TRN in the text). Kymographs were made using the ImageJ Kymograph plugin ^59^ and anterograde and retrograde movements were determined using a displacement of 2 µm as the minimum cut-off. Each mitochondrion was counted only once. The strains used in this study have been mentioned in Table S1.

### UNC-116::GFP and KLC-2::GFP imaging

L1 and L4 animals were anaesthetized with 30 mM Sodium azide and imaged using 60X/100X objective on a Olympus IX83 microscope with spinning disc (Perkin Elmer Ultraview, Waltham, MA, USA) with a Hamamatsu (SZK, Japan) monochrome EMCCD camera. Images for a specific cell type were collected from all genotypes grown identically, at the same time, using identical imaging conditions. Imaging conditions varied for different cell types.

### Western blotting

Protein extraction using Homogenization buffer (HEPES pH7.6-15mM; KCl-10mM; MgCl_2_-1.5mM; EDTA-0.1mM; EGTA-0.5mM; Sucrose-44mM in water) with Protease Inhibitor Cocktail (Complete Mini EDTA-free Protease Inhibitor Cocktail Tablets; Roche; Catalog No. 04 693 159 001) was carried out from a growing plate. Western blot analysis was carried out as described in ^60^. Polyclonal rabbit antibodies against UNC-116 and KLC-2 were raised by Bioklone, Chennai using HIS tagged proteins made from constructs as described earlier ^33^. We validated the antibodies on Western blot using protein extracts from *Ex[*_*p*_*unc-116::unc-116::gfp]; unc-116(f122)* and *Ex[*_*P*_*klc-2::klc-2::gfp]; klc-2(km28)* worms (data not shown). Antibody dilutions used were Rabbit α-UNC-116 (1:40000), Rabbit α-KLC-2 (1:40000), mouse α-Actin (1:1500) (NeoLabMarkers)]. Secondary antibodies dilutions were anti-Rabbit-HRP (1:10000), anti-Mouse-HRP (1:6000) (ABclonal). Three or more independent biological repeats were carried out in duplicates.

### Quantitation of levels of UNC-116 RNA

RNA was extracted from whole worm lysate from a just starved plate using Sigma TRI reagent. cDNA was synthesized using Superscript IV Reverse Transcriptase (ThermoFisher Scientific) using random hexamers. qPCR was done with Kapa SYBR FAST universal qPCR mastermix on Roche LightCycler-LC96. For purposes of quantitation, the samples were normalized to wild type. At least three independent biological repeats were carried out with at least two technical repeats. No genomic DNA contamination was observed when using actin primers. Quantitation was carried out as described in ^61^.

### Generation of domain structures for Miro

The domain structure has been generated using ScanProsite ^62^ and MyDomains ^63^. The genomic DNA sequences were sourced from the NCBI Database.

### Primers sequences used

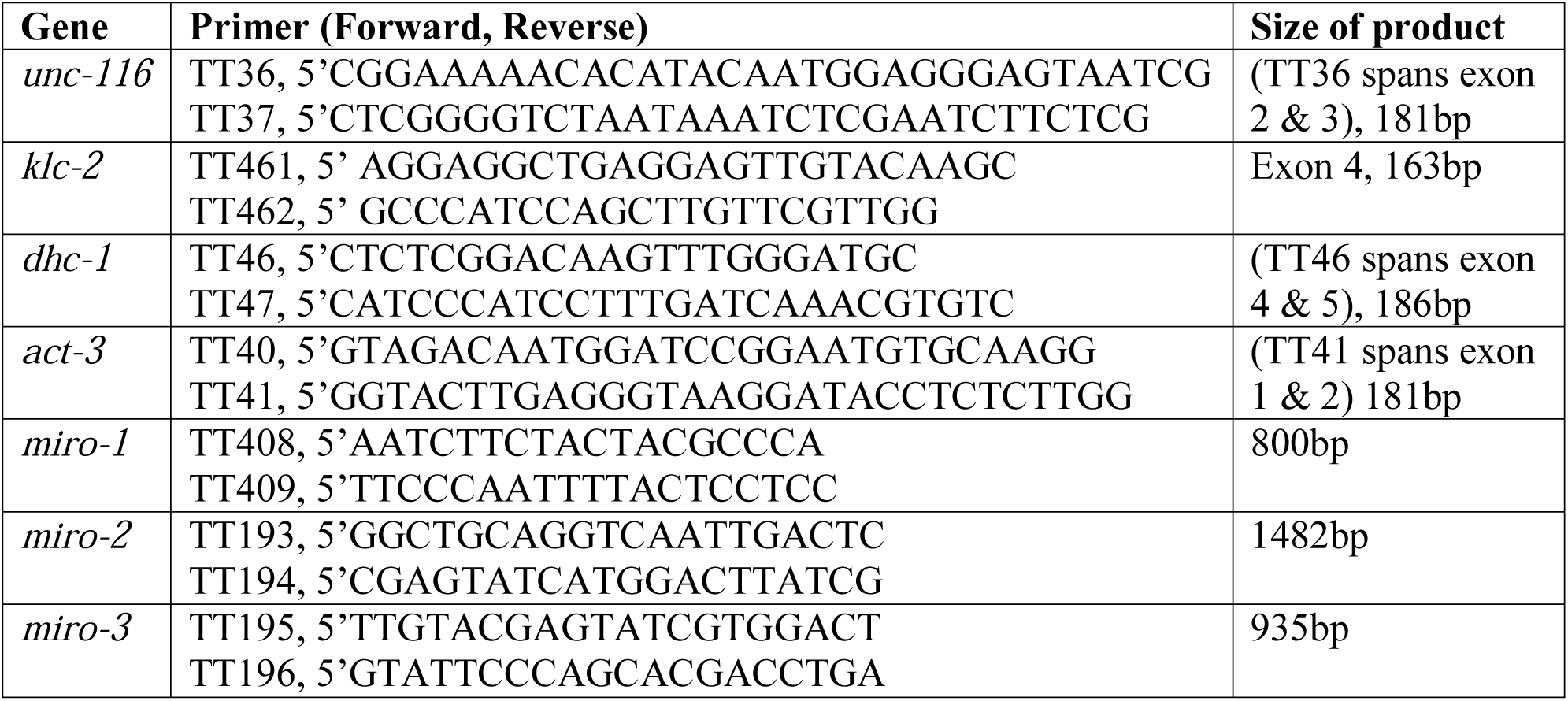

## Data availability

All data and reagents are available upon request.

## Author contributions statements

Designed research: Sandhya P Koushika (SPK), Guruprasada Reddy Sure (GPRS), Anusheela

Chatterjee (AC), Nikhil Mishra (NM), Vidur Sabharwal (VS)

Conducted research: All authors

Interpreted data: SPK, GPRS, AC, NM, VS

Wrote the paper: SPK, AC, NM

## Acknowledgements

This paper is dedicated to the memory of GPRS who passed away in April 2013 in a motor cycle accident. *jsIs609* was made by SPK in Dr. M. Nonet’s laboratory. We thank Drs. Yishi Jin, Francis McNally and Kenneth Miller for reagents. We thank Drs. Krishnamurthy, Manoj Matthew at CIFF, NCBS supported by the DST – Centre for Nanotechnology (No. SR/55/NM-36-2005). We thank Parul Sood and Neena Ratnakaran for confocal images. We thank Souvik Modi for discussions and help during analyses of *miro* domains. We thank Shubha Shanbhag for integrating UNC-116::GFP and KLC-2::GFP transgenic lines. We thank Salik Ansari for help in building strains. Some strains were provided by the CGC, which is funded by the NIH Office of Research Infrastructure Programs (P40 OD010440). *miro-1(tm1966), miro-2(tm2933)* and *miro-3(tm3150)* was provided by the Mitani Lab through the National Bio-Resource Project of the MEXT, Japan. This study was supported by grants to SPK from DBT, DBT overseas associateship, HHMI and TIFR. Salary support comes from TIFR and HHMI (NM), TIFR (AC, NM), CSIR (GPRS), UGC (SD), BITS (AA) and DST (SM). The spinning disc confocal microscope was supported by HHMI and DAE Project 12-R&D-IMS-5.02-0202. We thank members of the SPK lab for their critical input on the manuscript.

