## Supplementary Materials for "UNC-16/JIP3 and UNC-76/FEZ1 limit the density of mitochondria in *C. elegans* neurons by maintaining the balance of anterograde and retrograde mitochondrial transport"

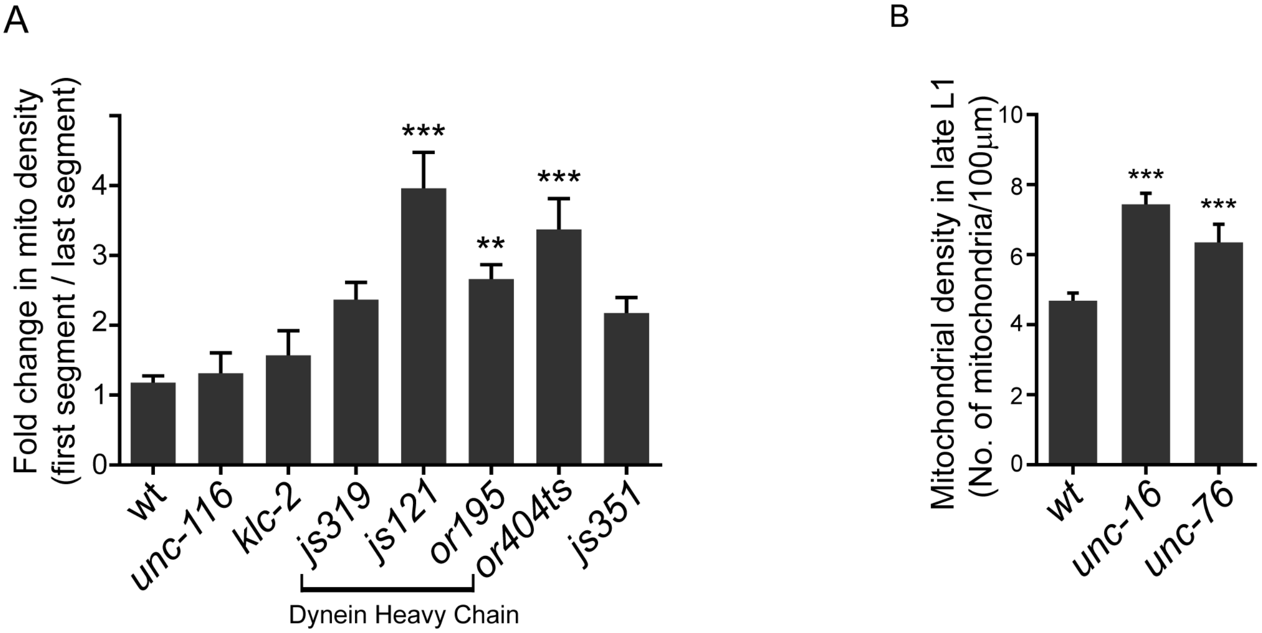


**Figure S1:** **(A)** Fold change in mitochondrial density between the first and last segment in wild type (*jsIs609*), *unc-116(e2310), klc-2(km11), dhc-1(js319), dhc-1(js121), dhc-1(or195), dnc-1(or404ts)* grown at 22$^{O}$C and *dli-1(js351)*. n≥7 worms. All comparisons made to wild type. Comparisons that are not significant are not listed. **(B)** Average density of mitochondria in late L1 animals of wild type (*jsIs609*), *unc-16(tb109)* and *unc-76(n2397)*. n≥19 worms. Data represented as Mean ± SEM. Statistical tests one-way ANOVA with Bonferonni multiple comparisons correction, p value **<0.01, ***<0.001.

A

```
Miro3      MPKVYSFRAFLQVKYGDREIFTDCSIN---NKSISK-IKLFGIVRVSWTYRLRKKMKI--
Miro2      ------------------------------------------------------------
Miro1      --------------MSDDETLADVRIVLIGDEGCGKTSLVMSLLEDEWVDAVPRRLDRVL

Miro3      ----------------------GLCWKS--ARRMSFVSSTRLQTSPLLMEFKQSTWLPLI
Miro2      --------------MDLSIKEEDENWIVSEIRQANVICVVYSVTDESTVDGIQTKWLPLI
Miro1      IPADVTPENVTTSIVDLSIKEEDENWIVSEIRQANVICVVYSVTDESTVDGIQTKWLPLI
                                 .  *     *: ..:. .   *.   ::  *:.*****

Miro3      RQSFGEYHKTPVILVGNKSDGTANNTDKILPIMEANTEVETCVECSARTMKNVSEIFYYA
Miro2      RQSFGEYHETPVILVGNKSDGTANNTDKILPIMEANTEVETCVECSARTMKNVSEIFYYA
Miro1      RQSFGEYHETPVILVGNKSDGTANNTDKILPIMEANTEVETCVECSARTMKNVSEIFYYA
           ********:***************************************************

Miro3      QKAVIYPTRPLYDADTKQLTDRARKALIRVFKICDRDNDGCL------------------
Miro2      QKAVIYPTRPLYDADTKQLTDRARKALIRVFKICDRDN----------------------
Miro1      QKAVIYPTRPLYDADTKQLTDRARKALIRVFKICDRDNDGYLSDTELNDFQKLCFGIPLT
           **************************************

Miro3      ------------------------------------------------------------
Miro2      ------------------------------------------------------------
Miro1      STALEDVKRAVSDGCPDGVANDSLMLAGFLYLHLLFIERGRHETTWAVLRKFGYETSLKL

Miro3      ------------------------------------------S-----------------
Miro2      --------------------------------------DGCLSPSELQNLFSVCPVSVIT
Miro1      SEDYLYPRITIPVGCSTELSPEGVQFVSALFEKYDEDKDGCLSPSELQNLFSVCPVPVIT
                                                     *

Miro3      --------------------------------------------------PSRAGNTLDS
Miro2      KD------------------------------------------------VGRAGNTLDS
Miro1      KDNILALETNQRGWLTYNGYMAYWNMTTLINLTQTFEQLAYLGFPVGRSGPGRAGNTLDS
                                                          .********

Miro3      IRVTRERKKDLENHGTDRKVFQCLVVGAKDAGKTVFMQSLAGRGMADVAQIGRRHSPFVI
Miro2      IRVTRERKKDLENHGTDRKVFQCLVVGAKDAGKTVFTQSLAGRGMADVAQIGRRHSPFVI
Miro1      IRVTRERKKDLENHGTDRKVFQCLVVGAKDAGKTVFMQSLAGRGMADVAQIGRRHSPFVI
           ************************************ ***********************

Miro3      NRVRVKEESKYLLLREVDVLSPQDALGSGETSAAVVAFLYDVSN--SFAFCATVYQKYFY
Miro2      NRVRVKEESKYLLLREVDVLSPQDALGSGETSADVVAFLYDVSNPDSFAFCATVYQKYFY
Miro1      NRVRVKEESKYLLLREVDVLSPQDALGSGETSADVVAFLYDISNPDSFAFCATVYQKYFY
           ********************************* *******:**  **************

Miro3      RTKTPCVMIATKVEREEVDQRWEVPPEEFCRQFELPKPIKFSTGNIGQSSSPIFEQLAMM
Miro2      RTKTPCVMIATKVEREEVDQRWEVPPKEFCRQFELPKPIKFSTGNIGQSSSPIFEQLAMM
Miro1      RTKTPCVMIATKVEREEVDQRWEVPPEEFCRQFELPKPIKFSTGNIGQSSSPIFEQLAMM
           **************************:*********************************
Miro3      AVYPHLRRVFYLNDSNLLSKITFGAAIVALAGLQHQSLAV
Miro2      AVYPHLRRVFYLNDSNLLSKITFGAAIVALAGLQHQSLAV
Miro1      AVYPHLRRVFYLNDSNLLSKITFGAAIVALAGFLVLKNL-
           ********************************:   .
```

B

| Protein | Allele | Deletion | Protein Domains (numbers denote amino acids) |
|---|---|---|---|
| MIRO-1<br>ΔMIRO-1 | *miro-1*(wt)<br>*miro-1(tm1966)* | ---<br>408 bp deletion<br>3 bp insertion | 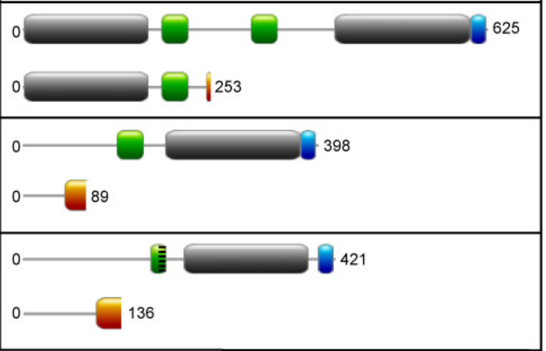 |
| MIRO-2<br>ΔMIRO-2 | *miro-2*(wt)<br>*miro-2(tm2933)* | ---<br>295 bp deletion<br>14 bp insertion | |
| MIRO-3<br>ΔMIRO-3 | *miro-3*(wt)<br>*miro-3(tm3150)* | ---<br>215 bp deletion | |



Keys: GTPase; EF hand; Partial EF hand; Transmembrane; Altered protein sequence

**Figure S2: (A)** Alignment of *C. elegans* MIRO-1, MIRO-2 and MIRO-3 proteins. **(B)** Protein domains in wild type and predicted protein domains in mutant *miro* alleles.

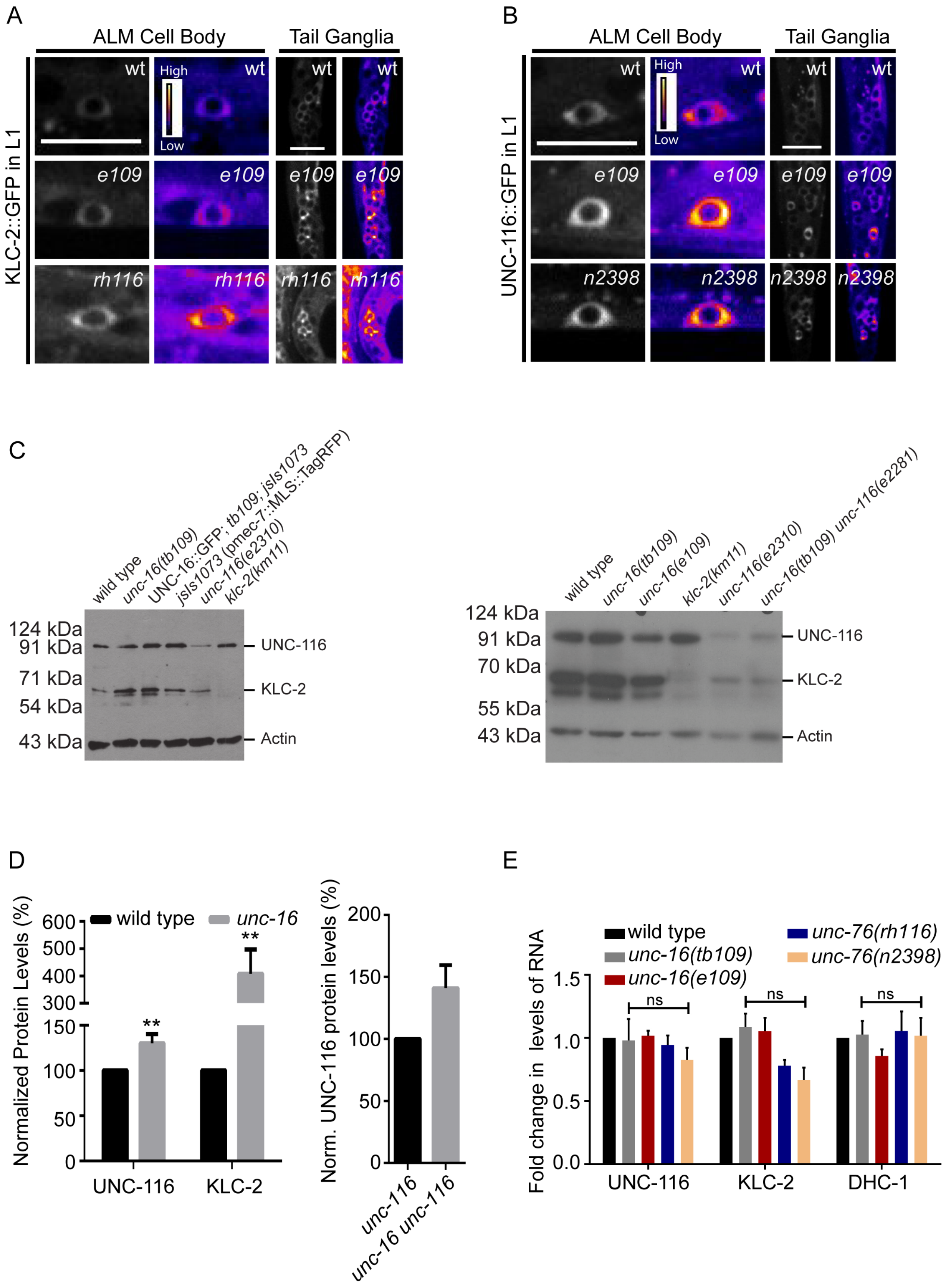
A
ALM Cell Body
Tail Ganglia
KLC-2::GFP in L1
wt
High
Low
e109
rh116
B
ALM Cell Body
Tail Ganglia
UNC-116::GFP in L1
wt
High
Low
e109
n2398
C
wild type
unc-16(tb109)
UNC-16::GFP; tb109; jsIs1073
jsIs1073 (pmec-7::MLS::TagRFP)
unc-116(e2310)
klc-2(km11)
124 kDa
91 kDa
71 kDa
54 kDa
43 kDa
UNC-116
KLC-2
Actin
wild type
unc-16(tb109)
unc-16(e109)
klc-2(km11)
unc-116(e2310)
unc-16(tb109) unc-116(e2281)
124 kDa
91 kDa
70 kDa
55 kDa
43 kDa
UNC-116
KLC-2
Actin
D
Normalized Protein Levels (%)
wild type
unc-16
**
UNC-116
KLC-2
Norm. UNC-116 protein levels (%)
unc-116
unc-16 unc-116
E
Fold change in levels of RNA
wild type
unc-16(tb109)
unc-16(e109)
unc-76(rh116)
unc-76(n2398)
ns
UNC-116
KLC-2
DHC-1

**Figure S3:** Levels of Kinesin-1 protein and RNA in *unc-16* and *unc-76* mutants.

**(A)** Representative images of KLC-2::GFP in ALM cell body and tail ganglia in L1 animals of wild type, *unc-16(e109), unc-76(rh116)*. **(B)** Representative images of UNC-116::GFP in ALM cell body and tail ganglia in L1 animals of wild type, *unc-16(e109), unc-76(n2398)*. **(C)** Full length blots showing levels of Kinesin-1 protein subunits. The observation with the following genotypes- (i) UNC-16::GFP; *unc-16(tb109); jsIs1073*, (ii) *jsIs1073* and (iii) *unc-16(e109)*- have not been analyzed for this study. These full length blots relate to Figure 3H. **(D)** Average normalized (norm.) intensity of Western blot signal showing elevated levels of both Kinesin-1 protein subunits in whole worm lysates prepared from *unc-16(tb109)* animals and UNC-116 protein levels in whole worm lysates of *unc-116(e2310)* and *unc-16(tb109) unc-116(e2281)*. *Unc-116(e2310)* and *unc-116(e2281)* are the same allele (see Table S1). All comparisons made to wild type in first graph and *unc-116(e2310)* in the second graph. Students t test, n≥3 independent protein extracts, western blots from each extract done in duplicates. **p<0.01. Second graph p=0.06 **(E)** Fold change in mRNA levels of *unc-116*, *klc-2* and *dhc-1* in *unc-16* and *unc-76* mutants. All comparisons are not significant (ns) and made to wild type using one-way Anova with Bonferroni multiple comparison correction.

**Table S1:** Strain List

| Genotype | Reference |
|---|---|
| *N2* | 57 |
| *jsIs609: Is[*$_p$*mec-7::mls::gfp]* | 30 |
| *jsIs1073: Is[*$_p$*mec-7::mls::rfp]* | 48 |
| *unc-116(e2310)* | *unc-116(e2310)* and *unc-116(e2281)* are the same allele 32 |
| *unc-116(rh24sb79)* | 33 |
| *unc-116(f122)* | 34 |
| *klc-1(ok2609)* | International *C. elegans* Gene Knockout Consortium 35 |
| *klc-2(km11)* | 21 |
| *klc-2(km28)* | 21 |
| *unc-16(tb109)* | 45 |
| *unc-16(e109)* | 44 |
| *unc-16(n730)* | 44 |
| *unc-16(ce451)* | 24 |
| *unc-76(n2397)* | 27 |
| *unc-76(n2398)* | 27 |
| *unc-76(rh116)* | 27 |
| *unc-76(e911)* | 27, 46 |
| *unc-14(ju56)* | 21 |
| *unc-14(e57)* | 47 |
| *dhc-1(or195)* | 37 |
| *dhc-1(js319)* | 36 |
| *dhc-1(js121)* | 36 |
| *dli-1(js351)* | 36 |
| *dnc-1(or404ts)* | 36 |
| *miro-1(tm1966)* | NBRP |
| *miro-2(tm2933)* | NBRP |
| *miro-3(tm3150)* | NBRP |
| *tbIs224* | We integrated rescuing extrachromosomal arrays- *Ex[*$_p$*unc-116::unc-116::gfp]* 48 |
| *tbIs235* | We integrated rescuing extrachromosomal arrays- *Ex[*$_P$*klc-2::klc-2::gfp]* 48 |
